# Open Raman Microscopy (ORM): A Modular Hardware and Software Framework for Accessible Raman Imaging

**DOI:** 10.1101/2025.10.26.684570

**Authors:** Kevin T. Uning, Henry Brisebois, Conor C. Horgan, Magnus Jensen, Yue Yuan, Shiyue Liu, Elzbieta Stepula, Steven Vanuytsel, Vishal Kumar, Stephen Goldrick, Robert D. Knight, Martin A. B. Hedegaard, Michael B. Albro, Mads S. Bergholt, Michael R. Thomas

## Abstract

Raman microscopy is a label-free, non-destructive imaging tool for spatially resolved chemical fingerprinting. Its powerful ability to reveal molecular information has driven rapid growth in applications across fields as varied as materials science, environmental analysis, and biomedical research. Despite its versatility, the accessibility of Raman microscopy is limited by expensive commercial setups and the technical barriers faced by researchers attempting to build custom systems. Here, we introduce an Open Raman Microscopy (ORM) framework based on a readily accessible modular microscopy platform. The ORM platform provides configurations for both high-throughput imaging and confocal imaging. We developed a dedicated python-based control and acquisition software, the ORM-Integrated Raman and Imaging Software (ORM-IRIS) designed to accommodate modular integration and control of components, including the laser source, spectrometer, and translational stages. Implemented across three institutions we demonstrate the ORM platform for high-throughput imaging of articular cartilage tissue, confocal three-dimensional imaging of a zebrafish embryo, and imaging of gold colloid decorated surfaces for surface enhanced Raman spectroscopy. Together, this open-source hardware and software framework enhances the accessibility of Raman microscopy across an expanding range of scientific applications.

## Introduction

Raman spectroscopy captures the characteristic vibrational signatures of materials, cells, and tissues with spatial detail that can approach the diffraction limit. This label-free, inelastic light-scattering technique provides molecular-level contrast that complements conventional imaging modalities such as brightfield or fluorescence microscopy. This unique capability has led to a plethora of applications in material science [1], forensics [2], art [3] and biomedical applications for cells [4], tissues [5] and nanotechnology [6]. Although commercial Raman microscopes are widely available, their high purchase and maintenance costs make them inaccessible for many laboratories, especially those in resource-limited settings. Moreover, as commercial systems evolve, many manufacturers incorporate proprietary technologies, such as preprocessing algorithms and automation features, that restrict user customization and maintenance. This lack of flexibility poses challenges in academic and interdisciplinary research environments, where experimental adaptability and transparency are essential. Consequently, there is a growing demand for accessible, open, and modular Raman microscopy solutions that reduce financial and technical barriers while maintaining high performance and reproducibility.

Open-source microscopy has gained substantial traction in recent years, ranging from low-cost widefield microscopes and light sheet microscopy to super-resolution microscopes. In terms of hardware frameworks, open-microscopy frameworks have followed the open-hardware trend [7] with the emergence of simple, low cost and agile microscopes such as FlyPi [8], OpenFlexure [9], UC2 system [10], µCube [11] and Octopi [12]. These microscopes have catapulted advanced microscopy from a scarce resource to everyday tools of life scientists and hobby enthusiasts alike. Open-microscopy software projects, such as Pycro-Manager [13], Micro-Manager [14], and the OMERO [15] have helped to accelerate progress, providing flexible and generalizable frameworks for building microscope hardware and enabling their control. However, none of these software ecosystems are specifically designed to support the data-intensive, hyperspectral mapping workflows required by Raman microscopy. In particular, they lack native tools for handling large multidimensional spectral datasets and managing the complex acquisition strategies often needed for chemically specific imaging. A dedicated open resource for Raman microscopy would not only allow for its wider adoption but also, encourage multi-modal microscopy configurations that meet the requirements of countless applications.

There are several essential requirements for an open Raman microscopy initiative that must be addressed and guided by three key principles: build simplicity, component accessibility, and modularity. First, the system should be simple enough to replicate by users with limited experience in optomechanics. Second, all components should be sourced from widely available suppliers. Third, the design should support modular integration to enable customization and multi-modal imaging applications. In addition, the platform should be configurable to accommodate different imaging modes, ranging from high throughput large-area scanning to high-resolution, sub-micron analysis, thereby offering flexibility across diverse sample types and research needs. Although several custom-built Raman microscope designs have been reported [16–18], none satisfy all three of these criteria, and an open-source software framework for instrument control has yet to be developed. Such a framework should emphasize ease of use, modularity in instrument control, maintainability, and extensibility. The control software should be operable via an intuitive graphical user interface (GUI) without requiring prior technical experience. It should also accommodate instruments from multiple manufacturers, enabling users to implement hardware for specific Raman microscopy applications. Finally, the software must be designed for long-term maintainability and extensibility, allowing it to adapt to evolving community needs and changes in the programming ecosystem.

Here, we introduce Open Raman Microscopy (ORM) and ORM-Integrated Raman Imaging Software (ORM-IRIS) to lower barriers for adopting Raman microscopy. The ORM hardware framework, illustrated in Figure 1a, leverages the modular Thorlabs Cerna® platform, though comparable commercial products or custom-built alternatives can also be used, and supports components from a range of instrument suppliers, including spectrometers, lasers, stages, and cameras. The ORM framework supports a high-throughput (HT) configuration for rapid imaging or a confocal imaging configuration for higher spatial resolution (Figure 1b). To meet the requirements of an open-source software framework, ORM-IRIS was developed with several key design principles. First, ORM-IRIS includes built-in controllers for instruments from common manufacturers, enabling diverse hardware configurations. Second, the software is written in Python to ensure ease of development and long-term maintainability, allowing new functionalities, such as scanning modes, calibration routines, and analysis tools, to be rapidly implemented and integrated (Figure 1c). Third, ORM-IRIS features a flexible core architecture with a prebuilt “extension” platform that provides access to core functionalities for creating custom modules, macros, and tools (Figure 1d). Here we demonstrate ORM implementation across three institutions with slightly different modular configurations and case studies for high-throughput imaging of cartilage tissue, imaging of self-assembled monolayer on gold colloid decorated surfaces for surface enhanced Raman spectroscopy, and confocal three-dimensional imaging of a zebra fish. These results demonstrate how the ORM framework can empower the research community with an open, adaptable, and cost-effective platform for advancing the adoption of Raman microscopy. The hardware design and assembly instructions of ORM are freely available at http://www.openramanmicroscopy.org/ and the ORM_IRIS software is freely available through our GitHub page at https://github.com/The-Thomas-Lab-UCL/orm-iris.

**Figure 1.**
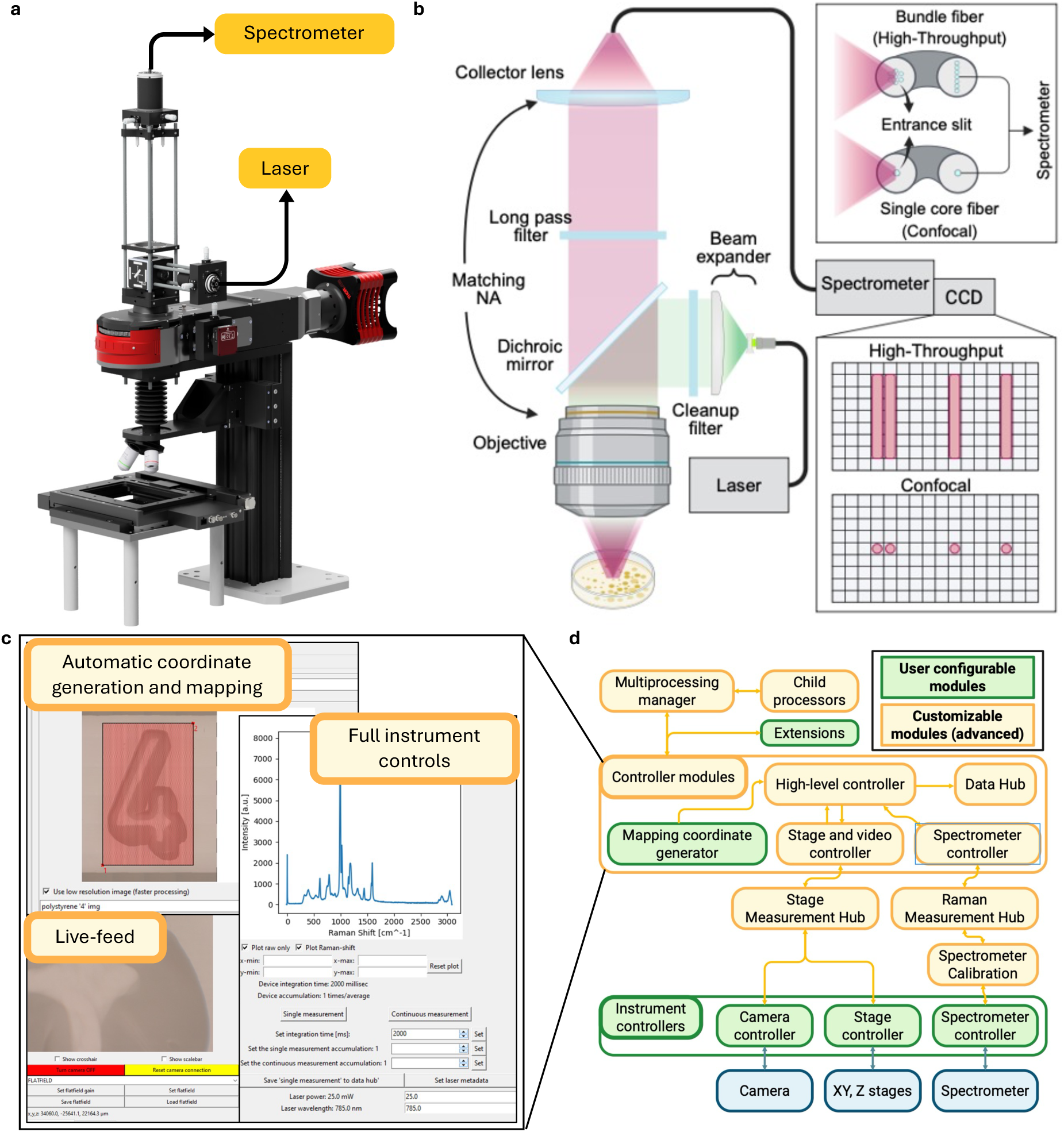
Overview of the ORM and ORM-IRIS. (a) 3D rendering of the ORM platform. (b) Diagram of the microscope optical design; the Raman signal collection can be switched between a round-to-linear fiber bundle for high throughput imaging or multimode fiber (for confocal imaging) (c) Graphical User Interface (GUI) of the ORM-IRIS capable of full instrument controls, live feed, automatic mapping, brightfield tiling, data management, and more. (d) ORM-IRIS software framework highlighting the modularity of the user configurable modules (green) and the advanced customizable modules (yellow), depending on the external devices (blue).

## Results and Discussion

### Open Raman Microscopy (ORM) Overview

The Open Raman Microscopy (ORM) platform presented here provides a flexible, high-performance framework for Raman imaging and spectroscopy. The system integrates down to diffraction-limited optical performance with modular hardware and open-source control software, enabling either rapid or high-resolution molecular imaging. Its architecture replicates the functionality of state-of-the-art confocal Raman systems while maintaining accessibility, adaptability, and cost efficiency for a wide range of research applications [16], [17], [18].

The microscope employs an epi-detection optical configuration (Figure 1a,b), where both excitation and collection occur through the same microscope objective. The platform accommodates multiple excitation sources through interchangeable laser modules and beam coupling using dichroic mirrors. The excitation laser is coupled via a single-mode fiber, beam expander and laser bandpass filter to fill the back aperture of the objective to ensure a diffraction limited clean Gaussian mode at the sample plane. The sample is mounted on a piezo or motorized (XY and Z) translation stage, providing automated raster scanning for Raman imaging with submicron precision. Stage scanning can be conducted in discrete sampling mode or in continuous sampling mode for faster scanning (*see Materials and Methods*). A dichroic beam splitter separates the backscattered Raman signal from the excitation path, directing the collected light through a long-pass edge filter to remove the intense Rayleigh-scattered photons while transmitting the weaker Raman-shifted signal. The filtered Raman scattered light is then coupled into the spectrometer via a single or bundled multimode fiber for wavelength dispersion and detection by a thermoelectrically cooled CCD array.

Two interchangeable detection approaches can be implemented depending on the specific application. In the confocal configuration, a single multimode (low-OH) silica fiber serves as the pinhole aperture. The effective pinhole size at the sample plane is determined by the fiber core diameter and the overall system magnification, which is set by the objective focal length and the tube lens. This size typically corresponds to ∼1-2 Airy units to balance signal throughput and axial optical sectioning. This arrangement enhances axial resolution by rejecting out-of-focus light and improving optical sectioning [19]. Alternatively, a round-to-linear (low-OH) silica fiber bundle can be used to increase collection efficiency for diffusely scattering samples such as biological tissues. In this configuration, the circular fiber input face samples a larger area in the microscope’s intermediate image plane, effectively relaxing the pinhole constraint (≫1 Airy unit) imposed by the objective/tube-lens magnification to maximize throughput at the expense of confocality. Inside the spectrometer, the output end of the bundle is rearranged into a linear array acting as a slit, effectively mapping the circular sampling region onto the CCD for vertical hardware binning. An important consideration is whether the linear array itself functions as a slit or if the spectrometer already includes an internal slit, as this determines both the spectral resolution and the optical throughput. Using no internal slit, this adaptable detection scheme allows the ORM platform to achieve very high collection efficiency and signal-to-noise ratio for diffusely scattering samples such as tissues. Together, the modularity establishes ORM as a powerful, open, and versatile tool for chemical imaging and spectroscopy across diverse materials and biological research contexts.

### Selecting matching lasers, filters, and spectrometer

Designing an effective Raman microscopy setup requires careful coordination between the excitation laser, optical filters, and spectrometer. Each component must be selected with consideration for wavelength compatibility, spectral resolution, and the intended sample type. The excitation laser defines both the achievable spectral range, obtainable spatial resolution, Raman signal intensity and the level of background fluorescence. Shorter wavelengths (e.g., 532 nm) provide stronger Raman scattering and higher spatial resolution but are prone to autofluorescence in some biological samples. Longer wavelengths (e.g., 633 nm or 785 nm) reduce fluorescence interference at the cost of weaker Raman signal intensity and spatial resolution. For most biological and soft-material applications, 785 nm lasers offer a good compromise between signal strength and autofluorescence suppression. In contrast, 532 nm is preferable for inorganic or cell samples where fluorescence is minimal or when high spatial resolution is required. Wavelength stabilized single mode lasers with narrow linewidths (<0.1 nm) are recommended to ensure spectral precision.

Optical filters must be matched precisely to the laser wavelength and to efficiently reject Rayleigh scatter while transmitting the Raman-shifted light. The ORM uses a typical configuration including a laser line clean-up filter in the excitation path and a long-pass edge filter in the collection path. For modularity, filters mounted in interchangeable holders simplify wavelength changes across laser configurations.

The spectrometer’s grating and detector must complement the chosen laser wavelength and desired spectral resolution. Grating efficiency peaks near specific blaze wavelengths; thus, selecting a grating optimized for the laser line maximizes signal. Typical groove densities range from 600–1200 grooves/mm, lower densities favor a wider spectral window and faster mapping, while higher densities improve spectral resolution for detailed chemical identification. A back-illuminated CCD or sCMOS detector with high quantum efficiency in the desired spectral range further enhances sensitivity and represents an important consideration.

The optimal system results from harmonizing these three components. A mismatch, such as a filter set not tuned to the laser line or a grating optimized for a different spectral region, can significantly degrade performance. The modular design of the ORM platform simplifies this process, allowing users to interchange lasers, filters, and spectrometers as required.

### System benchmarking

We benchmarked the imaging resolution and throughput of the developed ORM system using both diffusely scattering and reflective samples. Table 1 summarizes the optical resolution obtained for the two imaging modes using the chromium/silicon interface of an electron microscopy calibration target, as well as the Raman signal intensities measured from silicon and a highly scattering powdered aspirin tablet, acquired with a 785 nm laser using both a 10× and 40× objective (*see Materials and Methods*). The confocal configuration exhibited improved axial (Z) resolution compared to the high-throughput mode, while the lateral (XY) resolution remained similar, as it is primarily determined by the laser spot size for flat silicon. Since a 105 µm fiber was used as a pinhole, diffraction limited performance cannot be expected in this case although alternative fibers can be employed by users to meet their needs. The intensity data shows a pronounced increase in Raman signal from the diffusely scattering aspirin sample in the high-throughput configuration for both objectives, whereas the intensity from silicon remained comparable across configurations. The magnitude of signal enhancement for diffusely scattering samples depends on the sample’s optical properties including absorption, scattering and anisotropy coefficients. Overall, the high-throughput mode enables faster acquisition of scattering samples, albeit at the cost of reduced axial and XY resolution relative to the confocal configuration that will be sample dependent.

**Table 1.** Optical resolution comparison between the high throughput HT (7x 105 μm bundle-to-linear fiber paired with fiber collimator) and the high-resolution confocal (105 μm fiber with lens tube) mode of the standard ORM setup obtained using an MCS-1TR-XY electron microscopy calibration grid and collecting line profiles across the chromium-silicon features. Intensities are the measured Raman intensity, averaged over 100 measurements of a powdered aspirin tablet as an exemplary scattering sample, and the silicon region of the calibration target at 1595 cm^-1^ (C=O stretch) and 520 cm^-1^ (c-Si) respectively. Acquisitions were performed with a 785 nm laser at 45 mW with a 500 ms integration time using either a 10x Olympus Plan N or 40x Olympus UPlanSApo objective (NA 0.25 and 0.95 respectively).

| **Configuration** | **XY-Resolution**<br>**(Silicon)** | **Z-Resolution**<br>**(Silicon)** | **Intensity**<br>**(Powdered**<br>**aspirin tablet)** | **Intensity**<br>**(Silicon)** |
| --- | --- | --- | --- | --- |
| *HT*<br>(10x NA 0.25) | 10 $\mu\text{m}$ | 231 $\mu\text{m}$ | 2119 [a.u.] | 899 [a.u.] |
| *Confocal*<br>(10x NA 0.25) | 10 $\mu\text{m}$ | 50 $\mu\text{m}$ | 190 [a.u.] | 122 [a.u.] |
| *HT*<br>(40x NA 0.95) | 2.5 $\mu\text{m}$ | 24 $\mu\text{m}$ | 3042 [a.u.] | 270 [a.u.] |
| *Confocal*<br>(40x NA 0.95) | 2.5 $\mu\text{m}$ | 10 $\mu\text{m}$ | 142 [a.u.] | 137 [a.u.] |

### ORM Case Study 1: Molecular imaging of cartilage tissues

Raman spectroscopy offers a highly unique approach to molecular imaging of cartilage [5], [20], [21]. The mechanical integrity and functional performance of cartilage depend on the spatial organization and relative concentrations of predominant extracellular matrix (ECM) components, collagen (COL), glycosaminoglycans (GAGs), and water (H₂O). The ability to resolve these constituents with chemical specificity is therefore central to understanding tissue biomechanics, degradation, and repair processes. Figure 2a-h demonstrates the ORMs ability to characterize the ECM spatial distribution throughout bovine cartilage sections (full microscope configuration has been detailed in the Supporting Information). Multivariate regression model fits to the cartilage fingerprint Raman spectra using a library dataset of collagen, GAG and water accounted for on average 82% +/- 3% of the variation of the fingerprint spectra, enabling the generation high resolution images of the spatial distribution of Raman ECM constituent scores (GAG_score_, COL_score_, H_2_O_score_) (Figure 2e-f). Well known spectral features of connective tissues were observed at ∼856 cm⁻¹ (proline of collagen), ∼1300 cm⁻¹ (CH₂ twisting), ∼1450 cm⁻¹ (CH₂/CH₃ bending of collagen), and ∼1650 cm⁻¹ (amide I, C=O stretching of collagen). Analysis of high wavenumber spectral region enabled high resolution images of distribution of water-associated OH_area_ (Figure 2d). Raman images depicted a spatial distribution of GAG_score_ and COL_score_ through the articular cartilage depth topmost (1 mm), with GAG_score_ and COL_score_ increasing from superficial to middle to deep zones, consistent with prior characterizations of articular cartilage biochemical heterogeneities [22], [23], [24]. Importantly, this setup was able to capture the ECM distribution of the chondrocyte pericellular matrix, thus achieving comparable spatial resolution to previous work [5]. Overall, these results confirm that ORM not only reproduces established Raman biochemical mappings of cartilage but also extends the imaging capabilities to a higher throughput. The system’s spatial fidelity makes it particularly valuable for studies investigating connective tissue degradation, tissue engineering and regenerative therapies, and degenerative disease models.

**Figure 2.**
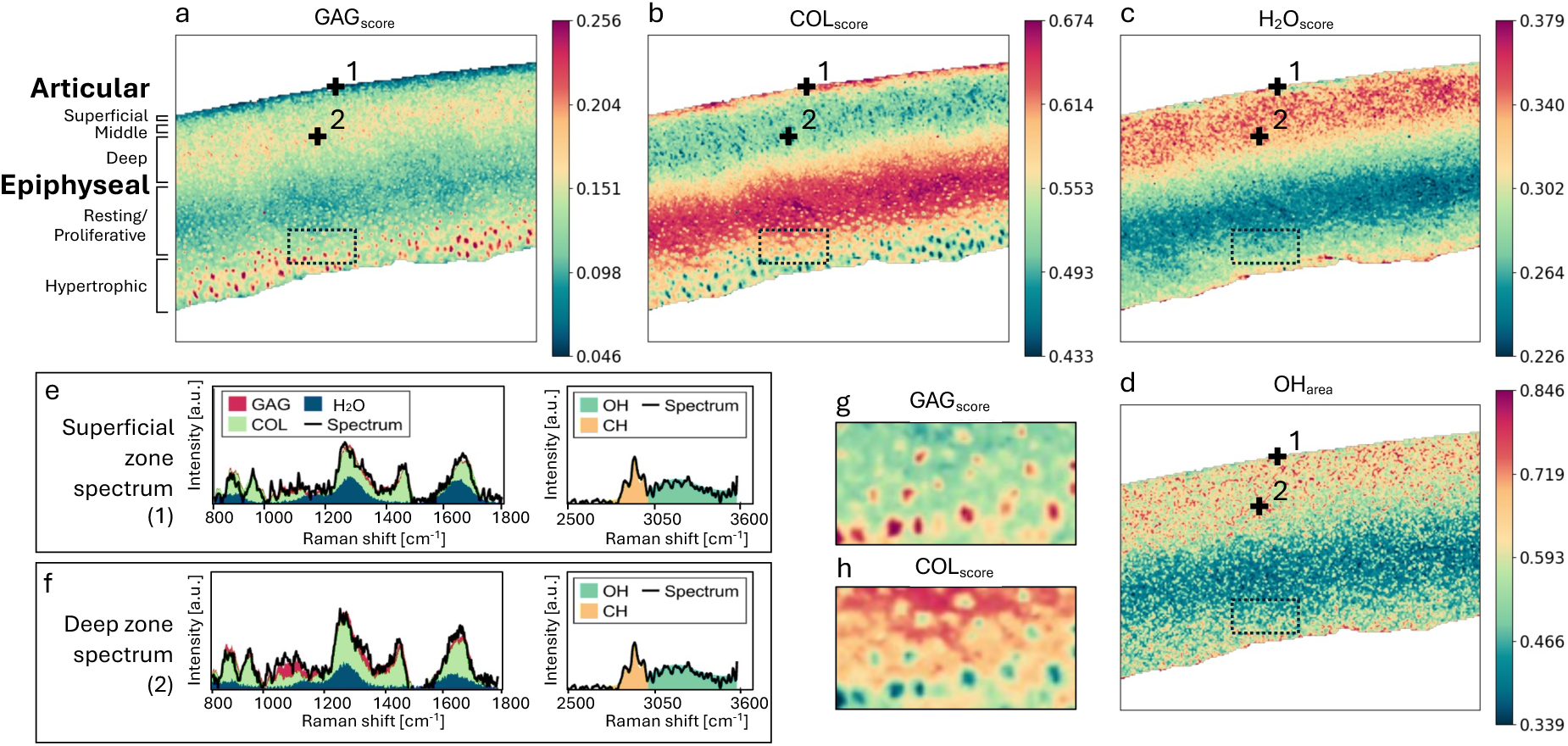
Raman spectroscopic imaging of bovine articular cartilage. (a–d) Multivariate Raman maps showing regression coefficients for glycosaminoglycans (GAG_score_), collagen (COL_score_), and water (H_2_O_score_) in the fingerprint region, and the water-associated OH bond peak area (OH_area_) from the high-wavenumber region. Scale bar: 500 µm. (e) Representative processed fingerprint Raman spectrum from the superficial zone, displayed as a stacked area plot showing cumulative contributions of COL, GAG, and H_2_O and a smoothed high-wavenumber spectrum showing area under the curve fitting for the CH and OH peaks. (f) Fingerprint and high-wavenumber spectrum from the deep zone. Zoomed-in GAG_score_ (g) and COL_score_ (h) Maps from the epiphyseal cartilage region, illustrating the ECM distribution around individual chondrocytes (location indicated by a broken black box in panels a–d). Scale bar 50 µm. The acquisition parameters were: 120 mW 785 nm laser with a 3 sec integration time, collected using a 1.0 NA 60x water-dipping objective. For high-throughput detection a 7x105 µm round to linear fiber bundle was used. The full setup optical configuration is available in Supporting Information.

### ORM Case Study 2: SERS substrate imaging

Surface-Enhanced Raman Spectroscopy (SERS) is a powerful extension of conventional Raman spectroscopy that enables ultrasensitive molecular detection by exploiting localized surface plasmon resonances generated by metallic nanostructures, typically composed of gold, silver, or copper. When molecules are positioned near these metallic nanostructures, the local electromagnetic field is greatly amplified by several orders of magnitude leading to a substantial increase in Raman scattering intensity. This plasmonic enhancement allows SERS to achieve single-molecule sensitivity in some cases and has led to widespread applications in chemical sensing, environmental monitoring, and biomedical diagnostics that have capitalized on the both the sensitivity enhancement of SERS but also the emergence of strongly localized effects at the nanostructure surface to establish a breadth of sensing strategies [25], [26], [27], [28], [29]. Despite its potential, SERS measurements require precise control over substrate fabrication and sample placement, making spatially resolved imaging a valuable tool for understanding and optimizing substrate performance [30]. Here we use the HT ORM configuration to demonstrate a scan of a SERS substrate comprising gold nanoparticles absorbed onto an aluminum surface shown in Figure 3 (a-b). Here the substrate was drop-cast with 4-mercaptobenzoic acid (MBA) and Rhodamine 6G (R6G) in discrete areas, dried and mapped. Unlike conventional Raman measurements, SERS signals are significantly enhanced where the MBA and R6G reside close to the gold nanoparticles and the hotspots between them. The measured data was processed following the protocol in the Materials and Methods section to reveal the SERS enhancement and the contribution of the MBA and R6G in each pixel of Figure 3 (a). Figure 3 (c-d) shows the Raman spectra acquired at identified points on the respective substrate maps for regions interfacing the regions of gold nanoparticles and those on aluminum. As a result of this enhancement SERS substrates typically facilitate reduced integration times during acquisition, shortening the overall scan time or facilitate more sensitive detection of trace analytes.

**Figure 3.**
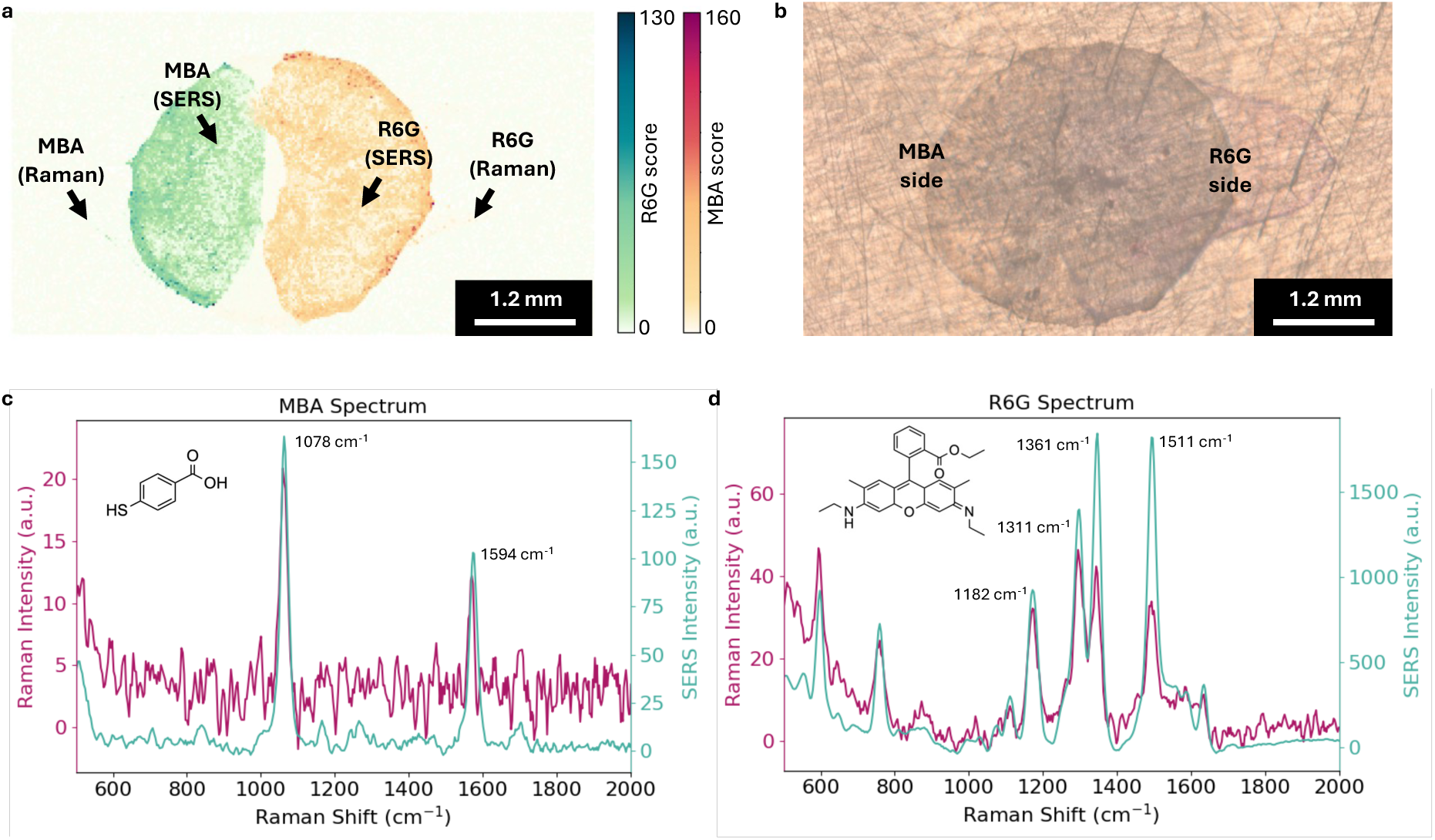
SERS substrate imaging. (a) False-color image of MBA and R6G deposited on a SERS substrate with MBA and R6G on its left and right-half respectively generated using a linear regression of the spectra obtained from point R6G(Raman) and MBA(Raman) indicates the contribution of MBA (green) and R6G (red) predominantly seen overlapping the location of gold nanoparticles shown in the brightfield image of the SERS substrate and drying patterns of R6G and MBA (b). (c) and (d) show Raman spectra obtained at each point identified in (a) either inside or outside the region containing gold nanoparticles. Acquisitions were performed with a 785 nm laser at 50 mW with 50 ms integration time, using the HT configuration with a 0.25 NA 10x objective.

### ORM Case Study 3: Volumetric 3D Raman imaging of zebrafish

We finally demonstrated use of the ORM system with a confocal configuration to perform volumetric Raman imaging of a zebrafish specimen (Figure 4a) [31]. The optical sectioning was critical for resolving fine structural features along the z-axis, enabling three-dimensional imaging at a resolution of 10 μm × 10 μm × 10 μm. This configuration allowed precise reconstruction of the fish morphology, capturing molecular signatures from anatomical sites such as the characteristic striped muscle pattern (Figure 4b). Principal Component Analysis (PCA) was applied to the Raman hyperspectral volume to identify the primary sources of spectral variation (Figure 4d–h). PC1 predominantly represented protein-rich tissues, while PC2 highlighted compositional contrasts between skin, muscle, and internal lipid-rich regions such as the yolk. Examination of PCA loadings and representative spectra revealed characteristic Raman peaks consistent with biological macromolecules. Distinct spectral features were observed at ∼1003 cm⁻¹ (phenylalanine ring breathing), ∼1300 cm⁻¹ (CH₂ twisting/lipid-associated plateau), ∼1450 cm⁻¹ (CH₂/CH₃ bending in proteins and lipids), and ∼1650 cm⁻¹ (amide I, C=O stretching of proteins). The 1450 cm⁻¹ and 1650 cm⁻¹ bands were particularly intense in muscle regions, indicative of the high protein and myofibrillar content, whereas the broader 1300 cm⁻¹ feature was more prominent in lipid-rich tissues of the yolk extension. Collectively, these results demonstrate that the ORM system can capture volumetric chemical information with high spatial and spectral fidelity. The 3D Raman reconstruction reveals distinct biochemical zonation within the zebrafish body, consistent with known tissue organization. Importantly, ORM achieves this without labels or staining, confirming its potential as a non-destructive tool for high-resolution, in situ molecular imaging of biological organisms.

**Figure 4.**
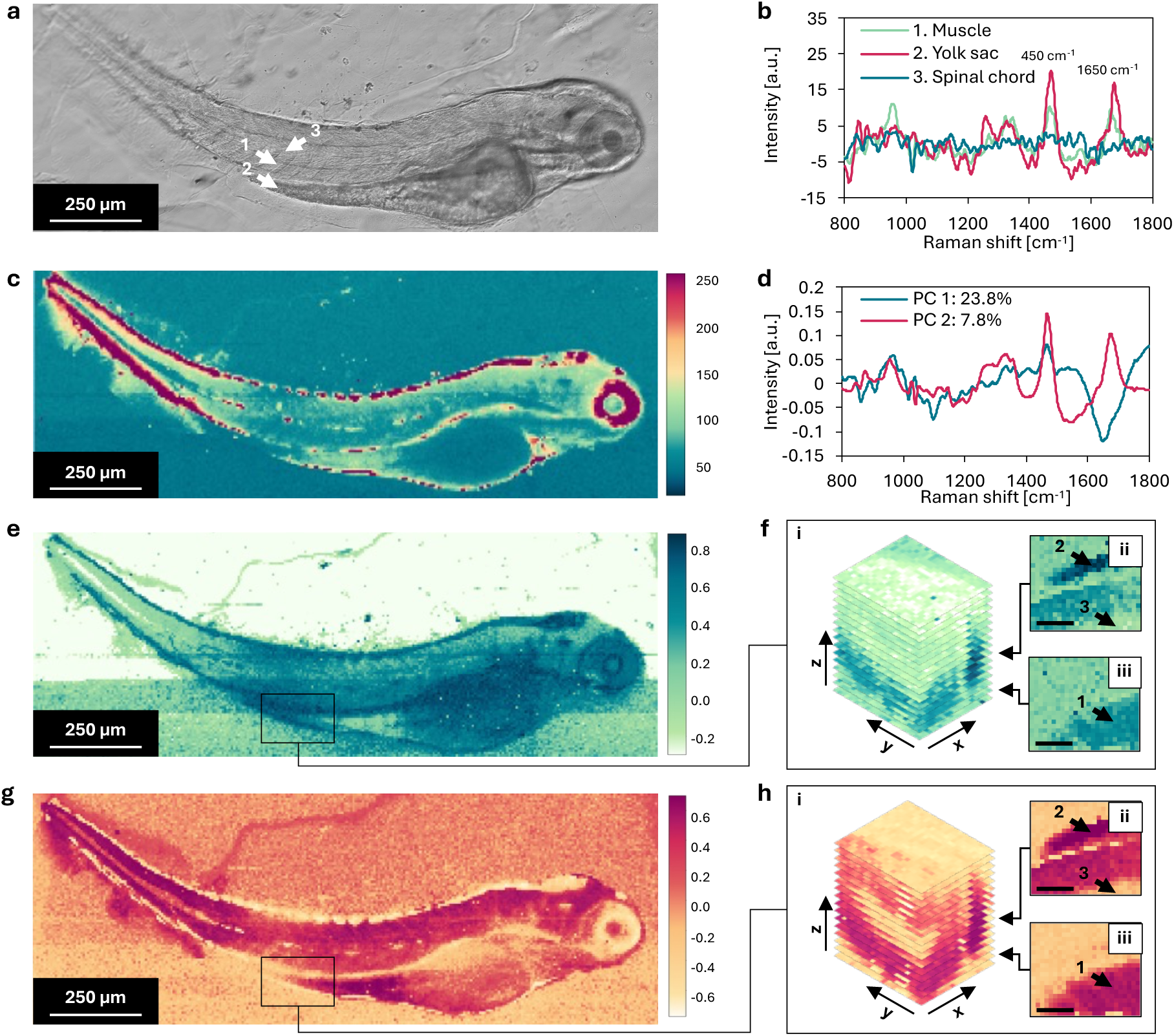
3D confocal Raman scan of a zebrafish. (a) Brightfield image of the zebrafish, the white arrows point to the 1. Fish muscle, 2. Yolk sack, and 3 Spinal chord, corresponding to the Raman spectra shown in (b) and to the black arrows in (f) and (g). The spectra show peaks of 1450 cm^-1^ and 1650 cm^-1^, due to the CH_2_ and Amide III bonds presence. Additionally, a small peak at 1004 cm^-1^ is also present representing phenylalanine, as well as the 1200 cm^-1^ to 1300 cm^-1^ representing Amide I. (c) Heatmap of the whole fish Raman scan at 1650 cm^-1^; the plot data is clipped at 250 [a.u.] such that the internal features of the fish are visible. (d) Pseudo spectra from the PCA analysis to plot the heatmap in (e-f) and (g-h) for the first and second vector respectively. The loading shows selectivity for spectra with signals around 1300 cm^-1^, 1450 cm^-1^ and 1650 cm^-1^. (e) 2D heatmap of the whole fish based on the first PCA component. The black rectangle shows the location of the 3D scan, with corresponding 2D slices shown in (f). (f.i) The full 3D stack of the slices, each separated by 10 μm, (f.ii) and (f.iii) 2D slice 70 μm and 20 μm from the bottom-most scan. The scale bars are 90 μm. (g-h) The same slices as in (e-f) but using the second PCA component. The acquisition was performed using a 785 nm laser, operating at 120 mW with a 1500 ms integration time and a 1.0 NA 60x water-dipping objective.

### Flexibility and customization

Multimodal integration represents an increasingly important strategy for expanding the analytical capabilities of optical microscopy platforms. While Raman imaging provides chemically specific information with minimal sample preparation, its relatively low scattering efficiency and longer acquisition times can limit spatial context. Integrating Raman microscopy with complementary imaging modalities can overcome these constraints by combining molecular specificity with structural, functional, or morphological contrast. In the ORM platform, brightfield or fluorescence imaging can serve as complementary channels that guide region-of-interest selection, enable rapid assessment of sample quality, and contextualize Raman spectra within morphological features. Brightfield imaging provides immediate visualization of tissue boundaries, cell distributions, or surface topography, while fluorescence can highlight specific structures based on molecular probes. Together, these modalities support more informed Raman acquisition strategies, reducing unnecessary mapping and improving throughput. Hardware integration is facilitated by shared optical paths, modular filter mounts, and coordinated stage control. By consolidating illumination, detection, and spatial positioning, multimodal workflows can be executed without requiring sample repositioning, thereby improving spatial registration between modalities. Software integration further streamlines data acquisition by synchronizing device control, image metadata, and hyperspectral mapping routines. This unified environment reduces operator overhead and enhances experimental reproducibility. Looking forward, the modular architecture of the ORM platform supports additional modalities such as polarization contrast, second-harmonic generation, or coherent Raman techniques. Incorporating these approaches could expand chemical sensitivity and enhance label-free contrast. Recent AI approaches also present an opportunity to enhanced throughput and data quality as photonic instrumentation and artificial intelligence converge [32], [33].

### Conclusions

In this work, we have introduced the Open Raman Microscopy (ORM) platform and its companion software, ORM-Integrated Raman Imaging Software (ORM-IRIS), as an open-source, modular, and accessible solution for high-performance Raman microscopy. The ORM framework combines diffraction-limited optical performance with flexible hardware and software architectures, enabling both high-throughput imaging for rapid sample screening and confocal volumetric imaging for high-resolution three-dimensional analysis. Through multi-institutional demonstrations, we showcased ORM’s versatility across a range of applications: high-throughput molecular mapping of cartilage sections, spatially resolved imaging of SERS substrates, and volumetric chemical imaging of zebrafish embryos. These studies highlight ORM’s ability to reproduce established Raman measurements while offering enhanced accessibility, configurability, and throughput. Importantly, the platform accommodates components from multiple suppliers and leverages Python-based software to enable modular integration, intuitive instrument control, and extensibility for future applications. By providing freely available hardware designs, assembly instructions, and open-source control software, ORM significantly lowers the technical and financial barriers to Raman microscopy, empowering researchers in academic, interdisciplinary, and low-resource settings. Collectively, these results demonstrate that ORM is a robust, adaptable, and cost-effective platform that can broaden the adoption of Raman spectroscopy across diverse fields, including materials science, chemical sensing, and biomedical research.

## Materials and Methods

### Open Raman Microscopy (ORM) hardware build

The microscope implementations described here are built around the Thorlabs Cerna® configuration which inherently facilitates modular component integration via standard optomechanics from various suppliers. ORM systems have been assembled using key components from common suppliers, including spectrometers and cameras from Ocean Optics, Wasatch Photonics, Andor (Oxford Instruments), and Princeton Instruments (Teledyne), as well as motorized stages from Zaber Technologies, Thorlabs, and Physik Instrumente. All these devices feature existing Python support or can be readily integrated due to the modular design of ORM-IRIS. The full list of the parts used to develop the ORMs is available in the supporting information, and the assembly instructions are available at http://www.openramanmicroscopy.org/. Briefly, the ORM consists of an epi-detection Cerna® microscope with a Raman module and brightfield imaging. To generate hyperspectral Raman images, motorized stages are used to move the sample in (XY) or (Z). Raman signal generation is achieved using a fiber-coupled excitation laser that enters the microscope system through a laser port, undergoes beam expansion, is cleaned up by a laser band pass filter. The laser light is coupled into the back aperture by a long-pass dichroic mirror and focused onto the sample. Scattered Raman signals are then collected by the objective and coupled through a long-pass filter to remove the Rayleigh scattered light. Finally, the Raman light is coupled into a multimode fiber (for confocal) or a fiber bundle (for high-throughput imaging). The fiber only acts as a confocal pinhole if a tube lens forms a real, magnified image of the sample onto the fiber face. Choosing the tube lens sets the magnification so the fiber core size can be matched with one Airy unit for confocal sampling, providing optimal lateral and axial resolution while rejecting out-of-focus background. Alternatively, one can instead couple with a simple collimator, where no image is formed, so the fiber no longer provides confocal filtering. Axial sectioning and resolution will degrade but the overall total collected signal will rise for samples depending on their optical properties. Where confocal operation is required, the tube lens configuration is employed, whereas if higher sensitivity is required, deeper penetration or if scattering samples are being measured, the bundle-to-fiber configuration can improve collection efficiency at a cost of spatial resolution as shown in Table 1. Brightfield imaging is obtained using the epi-illumination module’s filter turret by inserting a beam splitter to allow white light illumination from the rear of the epi-illumination module and image acquisition via a camera located on its side at an additional port.

### ORM-Integrated Raman Imaging Software (ORM-IRIS)

ORM-IRIS is a Python software dedicated to enabling the ORM hardware’s modularity and to provide an easy-to-use control. End-user configurable modules available include hardware controllers, coordinate generation methods, and the extension platform as highlighted in Supporting Figure 2. In addition to any new hardware controllers and coordinate methods, the extension platform allows users to develop custom functionalities, such as integrating pre-existing packages for real-time data analysis or macros to control the instrument operations. The ORM-IRIS GUI screenshot with a comprehensive interface, and its high-level structure are shown in Supporting Figures 5 and 6. The software framework is subdivided into three parts: (1) the core-functionality, (2) controller, and (3) support group. The core-functionality group consists of programming modules core to the app itself, such as the GUI, data-handler, and data processors. The controller groups consist of modules that bridge the communication between the instruments and the core functionality modules and make use of the Software Development Kit (SDK) provided by each respective manufacturer. This controller module separation is key in allowing facile modularity and the development of new instrument controllers. Finally, the support group consists of modules that support the operations of the core-functionality modules, such as the multiprocessing modules to parallelize tasks, and additional functionalities through the extension platform. Additionally, users can also create custom Python scripts for the different controllers and extensions, such as macros for the ORM-IRIS. It is open-source and is based on several packages to enable core functions: dill [34], keyboard [35], matplotlib [36], numpy [37], opencv_python [38], pandas [39], [40], Pillow [41], pyarrow [42], pyusb [43], scipy [44], and configupdater [45]. Instrument specific packages: zaber_motion [46], PIPython [47], pyserial [48], pylablib [49], and pythonnet [50]. The following Python-standard packages were also used: os, sys, glob, tkinter, threading, and multiprocessing. The ORM-IRIS is distributed under the GPL-3.0 license.

### ORM-IRIS: Spectrometer and objective calibration

Calibration is imperative in Raman and optical microscopy to ensure reproducibility. Specifically, the wavelength must be calibrated to an atomic emission lamp or a reference material, such as polystyrene [51], and similarly for the intensity, using reference material such as the NIST SRMs slides [52]. Once the reference table is obtained, ORM-IRIS can use it to automatically correct the spectrometer output intensity and wavelength, performed using polynomial regression for both [53]. An example calibration result is shown Supporting Figure 3 using polystyrene based on reference peaks [51].

Microscope objectives are calibrated so that pixel distances can be converted into physical units, enabling image corrections such as stretching and rotation, which are essential for functions like automated image tiling. The objective is calibrated by manually tracking a feature at different stage coordinates, which are used to calculate the x-scale [pixel/mm], y-scale [pixel/mm], image rotation [radians], and stage flip [Boolean]. These calibration results are stored locally for each microscope objective. Once calibration is completed, automated image tiling functionalities can be employed, where the user can select a region of interest (ROI), allowing the software to automatically acquire and tile images. The subsequent image can then be used to define a Raman mapping area and overlay the resulting Raman image.

### Stage control: discreet and continuous sampling

Raman imaging can be conducted using two sampling approaches: discrete and continuous mapping. In the discrete mode, the stage moves to each mapping coordinate, stops, acquires a Raman spectrum, and then proceeds to the next point. Because the system must wait for the stage to finish moving and settle at every position, this method is relatively time-consuming. Alternatively, Raman imaging can be conducted in continuous scan mode. In this mode, Raman measurements are collected while the stage remains in motion across the mapping coordinates. By eliminating the need to stop and wait at each point, this approach significantly reduces overall mapping time. In testing with a Zaber X-ASR100B120B-SE03D12 stage at a 500 ms integration time, mapping speed improved by up to a factor of five. Additional details are provided in the supporting information. One important consideration is that unlike the discreet mode that scans points, the continuous mapping mode integrates Raman signal across linear pixels. Therefore, any experimental impacts of this should therefore be considered by the user (e.g., such as photobleaching effects or sample inhomogeneity considerations).

### ORM-IRIS: Data format

Raman hyperspectral mapping data is multidimensional including timestamp, x-coordinate, y-coordinate, z-coordinate, wavelength and intensity data and requires an appropriate save-data structure. There exist currently no standardized data format. Hence, we developed our own format built around the SQLite3 database. It is structured such that multiple mapping measurements can be saved into one file, with the raw files stored externally with a parquet file type ensuring that data readability is independent of ORM-IRIS. Measurement data is saved in this format by default as it requires approximately 80% less disk space compared to .txt files (tested with 5×1000 data points Raman images which took 403 MB as txt files and 82 MB with the parquet structure). ORM-IRIS can convert these database files into other file types such as .txt files and .csv to ensure compatibility with the other software such as MATLAB or R. The save data structure has been designed in a such a way as to ensure support for alternative file types can be easily integrated if a standardized saving file structure is adopted by the evolving Raman microscopy field. In addition to the measured spectra and coordinates, the app also stores its metadata including the measurement timestamp of each obtained spectrum (measurement date accurate to microseconds), integration time, accumulation, laser wavelength, laser power, objective, acquisition mode (discreet/continuous), and hardware identifiers.

### ORM spatial resolution benchmarking

The spatial resolution was benchmarked using a TEM calibration grid, which has repeating lines of various spacings 1 μm, 2.5 μm, 5 μm, 10 μm, and larger. Specifically, the grid used was the MCS-1TR-XY traceable calibration standard from Labtech International Ltd., United Kingdom with chromium patterns on a monocrystalline silicon wafer. The XY resolution was benchmarked by scanning over the feature of known size and was tested with increasingly smaller features. The reported XY resolution is the smallest distinguishable spacing size of the grid, visualized at 520 cm^-1^. To benchmark the Z resolution, the silicon surface was incrementally moved through the focal plane while recording the Raman scattering. The intensity of the Raman peak at 520 cm⁻¹ was plotted as a function of the Z-position, and the Z resolution was determined from the Full Width at Half Maximum (FWHM) of this profile.

### Articular cartilage preparation

A full thickness cartilage explant (06 mm ξ 1.5 mm) was harvested from a 1-year-old bovine femoral groove. The explant was fixed overnight in a solution of 3.7% formaldehyde from Fisher Scientific (BP531-500), 5% acetic acid from Sigma-Aldrich (695092-500mL), and 70% ethanol Decon Laboratories, Inc. (2701). The explant was embedded in OCT and processed via cryostat into a 100 μm thick transverse section through the tissue depth. The section was affixed to a steel plate via a small drop of cyanoacrylate adhesive applied on the tissue lateral surface, away from the imaging region of interest, and submerged in phosphate buffered saline during imaging.

### Zebrafish preparation

Animals were reared at the Kings College London Zebrafish Facility and maintained in accordance with UK Home Office regulation, UK Animals (Scientific Procedures) Act 1986, under project license PP9727122. Embryos were obtained from natural spawning and embryonic fish were maintained in E3 Phenylthiourea (PTU) solution at 28.5°C to inhibit pigment formation medium [54]. Larvae were anaesthetized in 0.004% w/v Tricaine (Sigma-200 mg/ml) in E3 media and mounted in 1.5% w/v low melt agarose (Sigma) for imaging [55].

### Surface Enhanced Raman Spectroscopy (SERS) substrate preparation

The SERS substrate was prepared by dispensing 3 μL of 40 nm gold nanospheres (OD 20) from Nanocomposix (AUCR40-5M) onto a flat aluminum surface, and dried at 50°C for 5 minutes. Subsequently, 2 μL of a 100 μM aqueous solution of Rhodamine 6G (R6G) from Sigma Aldrich (R4127-5G) was dispensed on the right half of the substrate, dried at 50°C for 5 minutes, and then a 2 μL droplet of a 100 μM aqueous solution of 4-mercaptobenzoic acid (MBA) from Sigma Aldrich (706329-1G) was dispensed on the left half of the substrate and dried at 50°C for 5 minutes prior to imaging. Note MBA solution was initially dissolved to 1 mM in ethanol (200 proof ethanol, Fisher Scientific) and then diluted to 100 μM in DI water.

### Articular cartilage data processing

The Raman spectra were preprocessed according to the following protocol. First, the wavenumber was corrected by matching the characteristic peaks of a polystyrene reference spectrum. Cosmic ray artifacts were removed using a Savitzky-Golay filter (polynomial order 4, frame width 21) to smooth the spectrum. Absolute residuals between the raw and smoothed spectra were calculated, and points exceeding the mean plus three times the standard deviation were identified as outliers. These outlier intensities were replaced with the mean of neighbouring smoothed values within a ±5-pixel window. Following this, the background was subtracted from the spectra. Any negative intensity values originating from nonlinearities resulting from subtraction were set to zero. Next, the spectra were truncated into two regions of interests, the fingerprint region of ∼800 cm^-1^ to ∼1800 cm^-1^ and the high-wavenumber region from ∼2700 cm^-1^ to 3600 cm^-1^. The fluorescence signal in the fingerprint region spectra was then removed using a fourth-order polynomial fitting algorithm. Whereas the residual fluorescence background was modeled as an exponential decay, and fitted to the pre-CH_2_ peak region using nonlinear least-squares optimization. The fitted exponential background was then subtracted from the spectrum. Finally, the spectra were normalized by dividing spectra with the area under the curve. Hyaline cartilage is primarily composed of a type-II collagen (COL) fibril network, sulfated glycosaminoglycan (GAG) matrix and interstitial water. Therefore, the fingerprint spectra were fitted to a multivariate linear regression model:

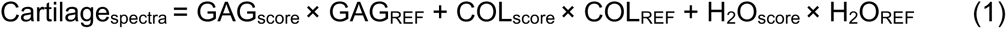

where: GAG_REF_, COL_REF_, and H_2_O_REF_, represent spectra of purified reference chemicals of each ECM constituent; the component “scores” are regression coefficients reflecting the relative contribution of each ECM constituent to the cartilage tissue spectra, as described in Kroupa et al. [56]. The reference spectra were obtained by measuring reference chemical spectra for water from UltraPure Distilled Water, Invitrogen (10977-015), collagen from Chicken Sternal Cartilage Type II, Signal-Aldrich (C9301), and GAG from Chondroitin Sulfate A Sodium Salt from Bovine Trachea, Sigma-Aldrich (C9819).

The high-wavenumber spectra were analyzed by measuring the areas under the carbon-hydrogen (CH_2_) and oxygen-hydrogen (OH) bond peaks, reflecting the organic and water content of the tissue, respectively, as described in Unal et al. [57]. A Savitzky–Golay filter was applied to the high-wavenumber spectra shown in Figure 2 (f) and (h) solely for visualization purposes; smoothing was not used in the quantitative analysis.

### SERS data processing

To calculate the contributions (also called scores) of MBA and R6G in the measured map, the data was first pre-processed as follows. First, the spectra were truncated from 300 cm^-1^ to 2100 cm^-1^ to remove the laser reflection below 300 cm^-1^ and the non-contributing region above 2100 cm^-1^ (for MBA and R6G). Second, a first-order Savitzky–Golay filter with a window of 3 was applied to remove any cosmic-rays. Next, a third-order polynomial regression was used to calculate and remove the background. Finally, this spectrum was fitted with a multivariate linear regression model:

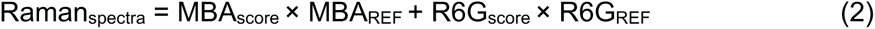

The reference signals were obtained from the same map of Figure 3 (a), from points outside of the SERS substrate, indicated by the MBA (Raman) and R6G (Raman) points. Therefore, the MBA and R6G contributions represent the relative signal enhancement magnitudes of the SERS substrate at each pixel. Finally, these values were shown as a false color image with the positive axis assigned to the MBA contribution and the negative axis assigned to the R6G contribution.

### Zebrafish data processing

Raman spectra acquired from zebrafish samples were pre-processed prior to analysis to enhance signal quality and minimize spectral artifacts. The raw spectra were first clipped to the wavenumbers of interest from 800 cm^-1^ to 1800 cm^-1^. Next, the background was removed from the spectra using an average of ∼1600 spectra adjacent to the fish. Third, spectral smoothing was applied using a first-order Savitzky–Golay filter with a window of 3 to reduce high-frequency noise while preserving peak shapes. Noise removal was then performed using a moving average of window size 5; this smoothing was separated from the Savitzky-Golay filter as it better preserves the peak shapes. Subsequently, third-order polynomial background subtraction was performed to remove fluorescence and baseline contributions. Finally, each spectrum was vector-normalized to correct for intensity variations and enable direct comparison across samples.

Principal Component Analysis (PCA) was used to separate out the features of Figure 4 (e – h), which was performed on the 2D fish scan, taking each wavenumber as a feature to obtain Figure 4 (e, g). The PCA vectors from this process were the used to project the 3D scan spectra to obtain Figure 4 (f, h).

## Acknowledgements

MRT acknowledges financial support of the Royal Society RGS\R1\201427. KTU acknowledges EPSRC dtpsa23242741. MSB acknowledges the European Research Council (ERC) under the European Union’s Horizon 2020 research and innovation program (Grant agreement No. 802778). RDK acknowledges support from the Leverhulme Trust (RPG-2020-105). MBA acknowledges support from the National Institutes of Health (R01AR081393). ES would like to thank the Early Career Engagement Award from The National Centre for the Replacement, Refinement and Reduction of Animals in Research in UK. The authors also thank Matthew Banner, John-Paul Ayrton, and Chapman Ho of the Department of Biochemical Engineering UCL for helpful discussions.

