## Supporting Information for "Open Raman Microscopy (ORM): A Modular Hardware and Software Framework for Accessible Raman Imaging"

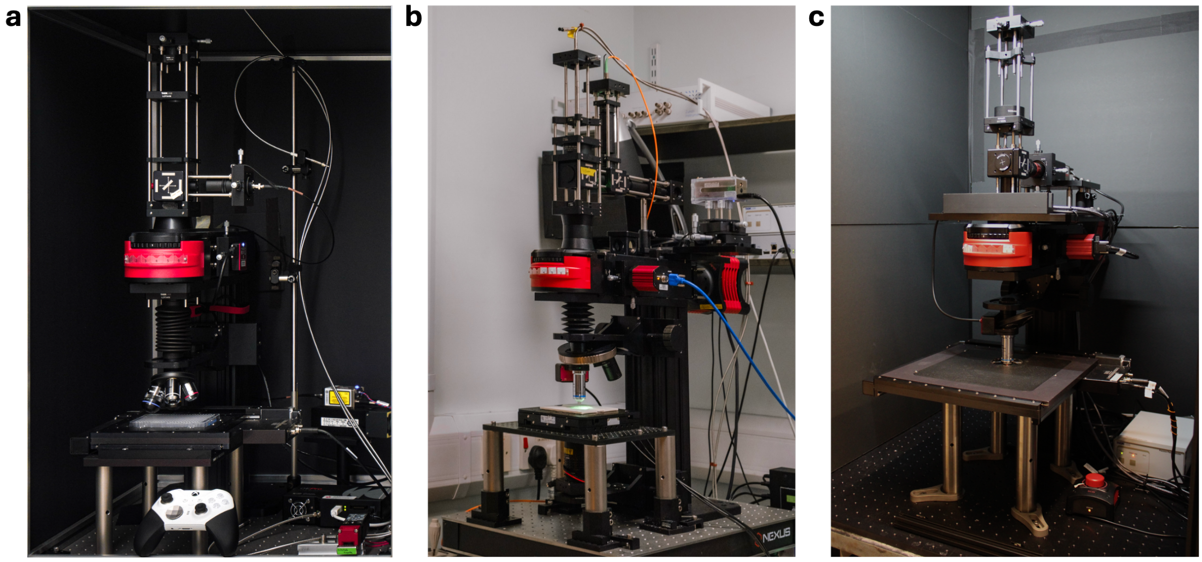

**Supporting Figure 1.** Photographs of two ORM example builds: (a) HT configuration uused in Case Study 1 with large travel range stages geared towards high-throughput measurements of standard 96 and 384 well plates. (b) Confocal configuration used in Case Study 3 configured for correlative fluorescence-Ramn imaging and piezo-stages geared towards high-resolution measurements. (c) HT configuration used in Case Study 2. HT makes use of a bundle-to-linear fiber for collection, and the confocal mode makes use of a single core fiber. These can be quickly swapped to switch between the confocal and HT modes when using the quick-fit magnetic tube lens adapter and performing appropriate alignment and calibration.

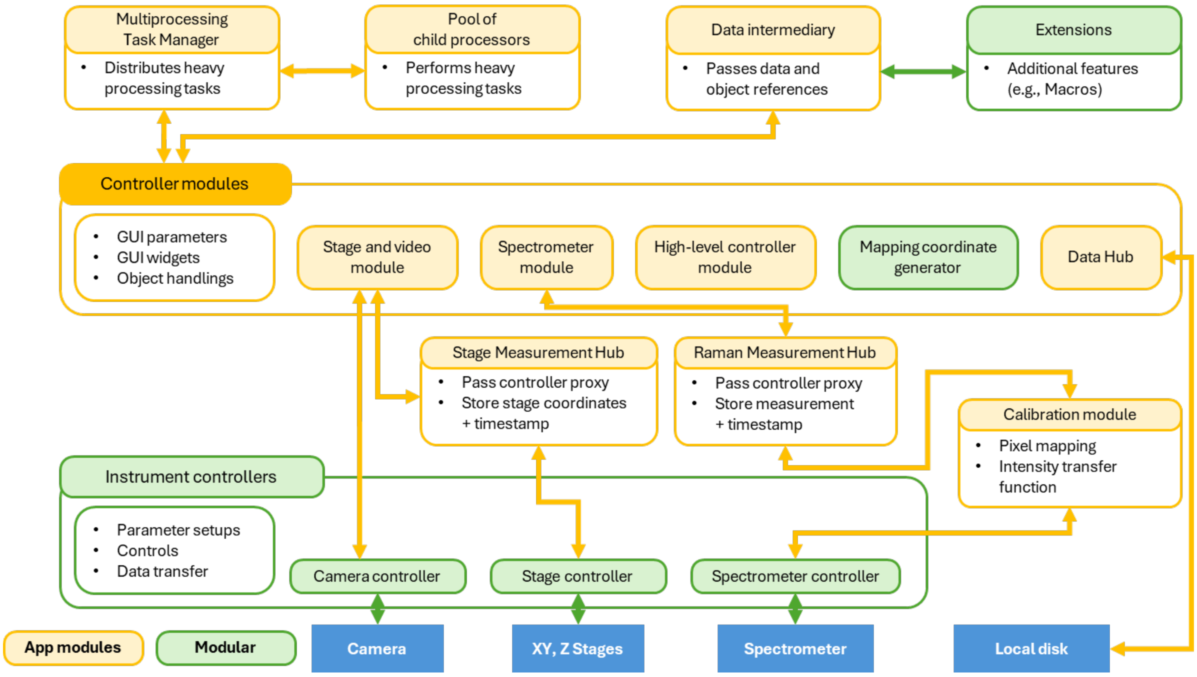


**Supporting Figure 2.** ORM-IRIS framework to facilitate ease of integration of new equipment and acquisition modules. The modules in [green] are user-replaceable to support different controllers, coordinate generation methods, and extensions. The rest of the modules are open for contributors to modify and update subject to review.

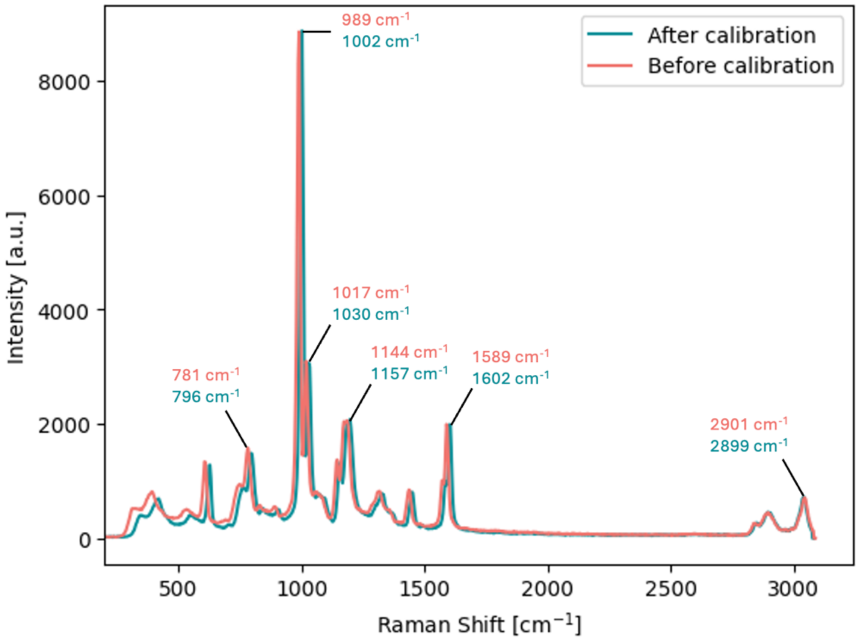


**Supporting Figure 3.** Reported polystyrene spectrum before and after a wavelength and intensity calibration. The calibration shifts the polystyrene spectrum according to the reference polystyrene peaks, as shown in Supporting Table 1.

**Supporting Table 1.** Polystyrene peak comparison between the reference, measurement prior to calibration and measurement after calibration using the ORM. The calibration was performed using a third order polynomial regression, which has brought the resulting measurement closer to the reference after calibration compared to before.

| Reference [$cm^{-1}$] | Measured before calibration [$cm^{-1}$] | Measured after calibration [$cm^{-1}$] |
|---|---|---|
| 795.8 | 781.49 | 796.1 |
| 1001.4 | 989.78 | 1001.99 |
| 1031.8 | 1017.77 | 1029.8 |
| 1155.3 | 1144.99 | 1156.53 |
| 1602.3 | 1589.13 | 1602.17 |
| 2904.5 | 2900.85 | 2898.31 |

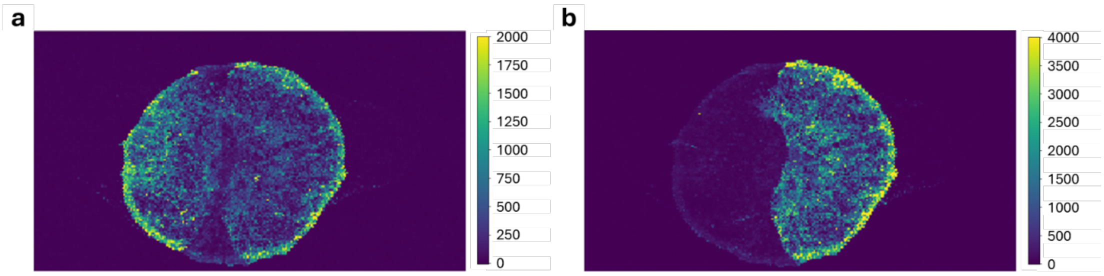


**Supporting Figure 4.** Raman maps of the SERS substrate shown in Figure 3 at: (a) the MBA peak of 1078 $cm^{-1}$ and (b) the R6G peak of 1361 $cm^{-1}$.

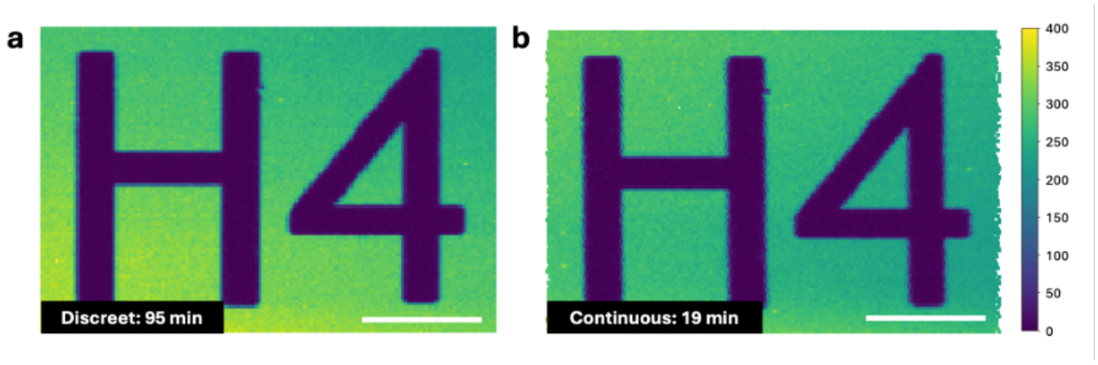


**Supporting Figure 5.** Raman mapping of an SEM finder grid ID at around 520 cm$^{-1}$ measured using: (a) discreet mode and (b) continuous mode, with 50 ms integration time, and 45 mW laser power. The discreet mode took around 95 minutes to finish, and the continuous mode took around 19 minutes to finish. The jagged edge on the left and right-side of the continuous map is caused by the inconsistent stopping distance inherent to the XY-stage and can be mitigated by obtaining a slightly larger scan area than required. The continuous mode is shown to be capable of achieving equivalent mapping result as the discreet mode with significantly less scan time required. The scalebar shows 20 μm.

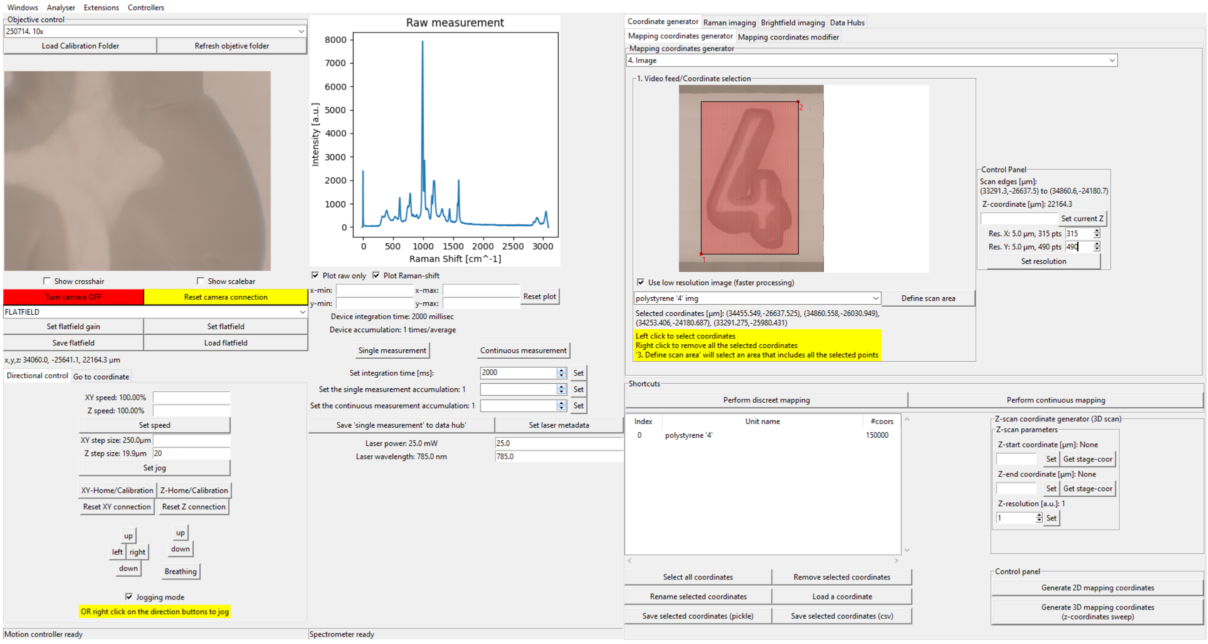


**Supporting Figure 1.** Example GUI screenshot of ORM-IRIS showing the XY-stage and camera control on the left, spectrometer control in the middle, and high-level controls on the right for: automatic (2D and 3D) coordinate generator and mapping, coordinate management system, brightfield tiling (hidden in a separate tab), data management (hidden).

### ORM Construction Details

The overall construction of the HT and Confocal ORM builds including cross-sections of key optomechanics can be seen in Supporting Figure 6. Here we describe in detail the components employed and in the following sections the components that differ from these in the builds used for generating Case Study 1 and 3.

### HT and Confocal ORM configurations (Table 1 & Case Study 2)

Motion control was achieved using a Zaber X-ASR100B120BSE03D12-KX14PG for the XY-stage and Thorlabs PLSZ with Thorlabs MCM301 Controller for the Z-stage. The laser used was an OBIS LX SF 785nm 50mW LASER SYSTEM:FIBER PIGTAIL FC/APC, guided via FC/APC collimator into a 2X beam-expander consisting of a Thorlabs N-SF11 Bi-Concave Lens, Ø25.4 mm, f = -25.0 mm, ARC: 650-1050 nm and a Thorlabs N-BK7 Plano-Convex Lens, Ø1", f = 50 mm, AR Coating: 650 - 1050 nm. The laser cleanup filter used was a Semrock 785 nm MaxLine® laser clean-up filter. The dichroic mirror used was a Semrock Di02-R785-25x36 785nm laser brightLine single-edge laser dichroic beamsplitter and the long-pass filter used was a Semrock 785 nm EdgeBasicTM best-value long-pass edge filter. The objectives used were the 10X Olympus Plan Achromat Objective, NA 0.25, 10.6 mm WD and a 40X Olympus UPlanSApo Objective, NA 0.95, 0.18 mm WD. A Thorlabs F810SMA-850 - Fiber Collimation Package, 850 nm, f = 36.20 mm, SMA collimator was used, coupling the collected light into the bundle end of a Thorlabs Linear Bundle, 1 X 7 Ø105/125 Fiber, Low OH, SMA, L=2m. This fiber was connected to a QE Pro Raman spectrometer from Ocean Optics employing a 100 µm slit. The HT configuration can be switched to the confocal configuration by switching the collimator with a Thorlabs AC508-180-AB-ML tube lens with a focal length matching that of the objectives (e.g 180 mm for Olympus) and the bundle fiber cable with a Thorlabs Ø105 µm, 0.22 NA, SMA905-SMA905 AR-Coated MM Patch Cable, 650 - 1100 nm, 2 m single core fiber optics cable. In theis build, the tube lens can be rapidly added or removed by using the Thorlabs LCP44F - 60 mm Removable Cage Plate magnetic holder to lower the switching time between the two configurations, taking less than 5 min to switch and re-calibrate the system.

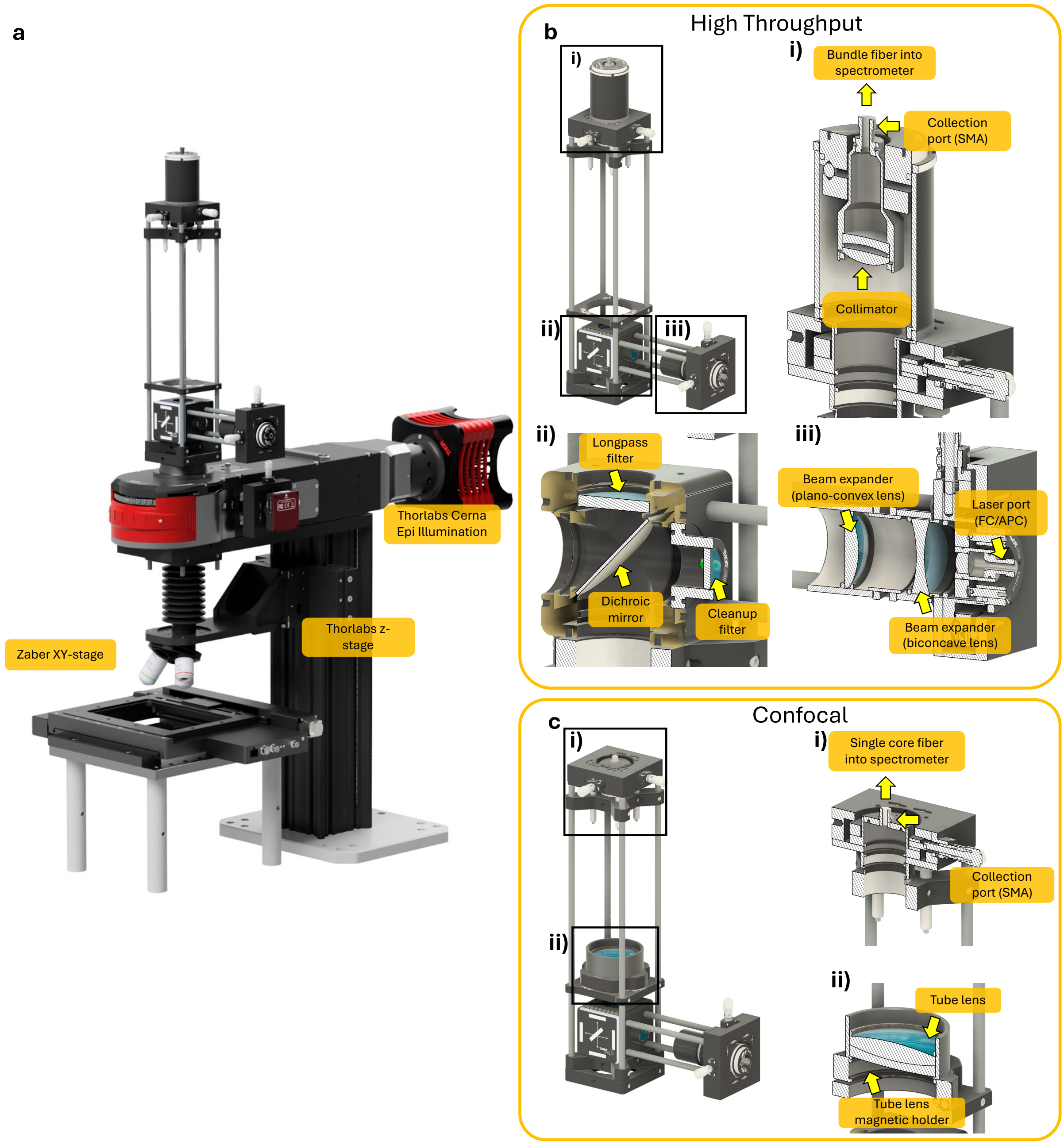


Supporting Figure 6. 3D renders with and without cross sections of (a) the ORM overall, and zoomed in views and cross-sections of (b) the HT and (c) Confocal configuration optomechanical constructions (employed in Table 1 and Case Study 2) that are mounted above the Thorlabs Cerna Epi Illumination system to better show the location of filters lenses and fiber collimators/ couplers.

#### ORM Configuration for Case Study 1: Cartilage scan

The cartilage scan was performed using an HT ORM configuration that employed the Thorlabs AC508-180-QB-ML sear coating 400-1100nm tube lens instead of a fiber collimator to guide light to the bundle-to-linear fiber to improve the spatial resolution while increasing the sensitivity compared to the confocal configuration. The main differences from the confocal build described above include a 785nm Toptica iBeam single mode diode laser that was used at 120 mW. The objective used was the 60x Nikon NIR Apo 1.0w, 1.0 NA, 2.8 mm WD. The spectrometer employed was an Ibsen photonics EAGLE Raman-S with an Andor iVac 316 Camera. The stages used were the Zaber X-ASR-E series motorized XY-stage and the Thorlabs PFM450 Z-stage.

#### ORM Configuration for Case Study 3: zebrafish scan

The zebrafish scan was performed using the same core confocal configuration described above. The main differences include a 785 nm Toptica iBeam single mode diode laser that was used at 120 mW. The objective used was the 60x Nikon NIR Apo 1.0w, 1.0 NA, 2.8 mm WD. The spectrometer employed was a QE Pro Raman spectrometer from Ocean Optics employing a 100 μm slit. The stages used were the piezoelectric Thorlabs PFM450E for the Z-stage and the piezoelectric U-751.24 stage with the C-867.2U2 controller, both obtained from Physik Instrumente (PI) SE & Co. KG.
